# Switching patterns of cortical-subcortical interaction in the human brain

**DOI:** 10.1101/2024.05.10.593351

**Authors:** Alessandro Nazzi, Chiara Favaretto, Antonino Vallesi, Maurizio Corbetta, Michele Allegra

**Affiliations:** Padova Neuroscience Center, University of Padova, Padova, Italy; Department of Neuroscience, University of Padova, Padova, Italy; Venetian Institute for Molecular Medicine, Padova, Italy; Department of Physics and Astronomy, University of Padova, Padova, Italy

## Abstract

Resting-state fMRI studies show that functional connectivity (FC) undergoes rapid fluctuations. Although the underlying neural mechanisms are poorly understood, a recent contribution analyzing stroke patients suggested that FC fluctuations involve a dynamic reconfiguration of cortico-subcortical interactions. Here, we analyze cortical-subcortical dynamic FC in a large cohort of healthy subjects (Human Connectome Project data base). Our analysis confirms that FC shifts are synchronized in cortex and subcortex. Two core subcortical ‘clusters’ comprising, respectively, limbic regions (hippocampus and amygdala) and subcortical nuclei (thalamus and basal ganglia) change their connectivity pattern with cortical regions. We consistently identify two recurring FC patterns (states). In state 1, limbic regions couple with the default mode network, in state 2 with sensorimotor networks. An opposite pattern is observed for thalamus/basal ganglia. Our findings hint at a general relevance of cortico-subcortical interactions in the generation of whole-brain spontaneous FC patterns.

## INTRODUCTION

Even in absence of external stimuli and overt behavior, the human brain thrives with activity^1,2^, most of which is sub-threshold and occurs in the infra-slow frequency range^3^. A striking feature of this spontaneous activity is that its fluctuations exhibit a well-defined spatiotemporal organization, with correlated fluctuations across the whole-brain that clearly emerge when looking at fMRI functional connectivity (FC) at rest. This phenomenon has been intensively scrutinized at the cortical level, leading to a well-established paradigm according to which activity fluctuations reflect the existence of several, canonical ‘intrinsic networks’^4^. In comparison, the subcortical level has received much less attention. Yet, subcortical structures may be fundamental for the generation of spontaneous activity, as cortical-subcortical loops are involved in most functional brain circuits according to a recent proposal^5^. Several studies have focused on single subcortical structures, such as the thalamus^6^, hippocampus^7^ or cerebellum^8^, analyzing their connectivity with the cortex and showing that they can be subdivided into regions associated with different cortical networks. An implicit assumption of these studies is that the functional coupling between subcortical structures (or subdivisions thereof) and the cortex is fixed, or ‘static’. However, FC at rest is time-varying, a phenomenon known as *dynamic functional connectivity*^9,10^ (dFC). While there is no universal agreement on the best methodology to characterize dFC, and on its meaning and causes^11^, evidence for dFC is abundant, as are results showing association between dFC and indicators of healthy and pathological cognition^12,13^. Therefore, the static picture of subcortical-cortical connectivity may hide a dynamic landscape where subcortical structures couple flexibly with cortical regions, and vice versa. Evidence in favor of this hypothesis comes from the recent study by Favaretto and colleagues^14^ (henceforth called FA22), who investigated the fluctuations in subcortical and cortico-subcortical FC in stroke patients. FA22 observed synchronized fluctuations in cortical and subcortical FC, and identified two main ‘blocks’ of highly synchronized subcortical structures (one comprising hippocampus and amygdala, the other thalamus and basal ganglia) that alternate between different patterns of connectivity with cortical networks. These findings suggested that cortico-subcortical interactions may be relevant for the dynamic reorganization of whole-brain FC, hinting at their general relevance in the emergence of whole-brain spontaneous activity patterns.

Here we build on FA22’s results and analyze dFC and cortico-subcortical interactions in a wide cohort of healthy young participants, made available by the Human Connectome Project^15^. The large sample size (N=1200) and fine temporal resolution (0.71 s) allow for a statistically reliable characterization of dFC, reducing instabilities due to individual variability, sampling variability, and artifacts. We wish to test several hypotheses, suggested by the FA22. First, we hypothesize the existence of synchronous connectivity shifts in cortex and subcortex, captured by different connectivity states or ‘dynamic functional states’. Second, we expect a split of subcortical structures into two groups characterized by internal synchronization and dynamic links with cortical networks. Third, in the light of previous work on the relationship between dFC and behavior, we conjecture that individual markers of cortical-subcortical dFC may contribute to explaining inter-individual variability in cognition. Finally, we assume that our findings will be robust with respect to details of the analysis pipeline used, including specific choices of cortical/subcortical parcellations.

## RESULTS

### Analysis overview

We considered a large sample of healthy human subjects (n=1078) from the Human Connectome Project^15,16^. For each subject, we extracted average BOLD time series for all regions of an atlas comprising 71 cortical regions and 19 subcortical and cerebellar regions from the Freesurfer atlas^18,19^. We computed functional connectivity (FC) matrices for sliding windows with a 60s duration, projected them onto the space spanned by the principal eigenvector, and vectorized them; we then concatenated together all time windows and subjects and performed a K-means clustering over windows. The resulting clusters are termed *dynamic functional states (DFSs)*.

In Fig. 1a we show the average BOLD signal of two cortical networks (sensorimotor and default mode) and two subcortical regions (thalamus and hippocampus). Starting from sliding-window functional connectivity, we extract the DFSs (Fig. 1b), which correspond to different cortical-subcortical interactions. This is evident in Fig. 1d, where we show dynamic variations of FC associated with the different DFSs. In DFS1, task-negative regions correlate positively with SC2 and negatively with SC1, while the opposite pattern is observed for task-positive regions. In DFS2, this trend is reversed, as task-negative regions correlate negatively with SC2 and positively with SC1, while the opposite pattern is observed for task-positive regions.

**Fig. 1.**
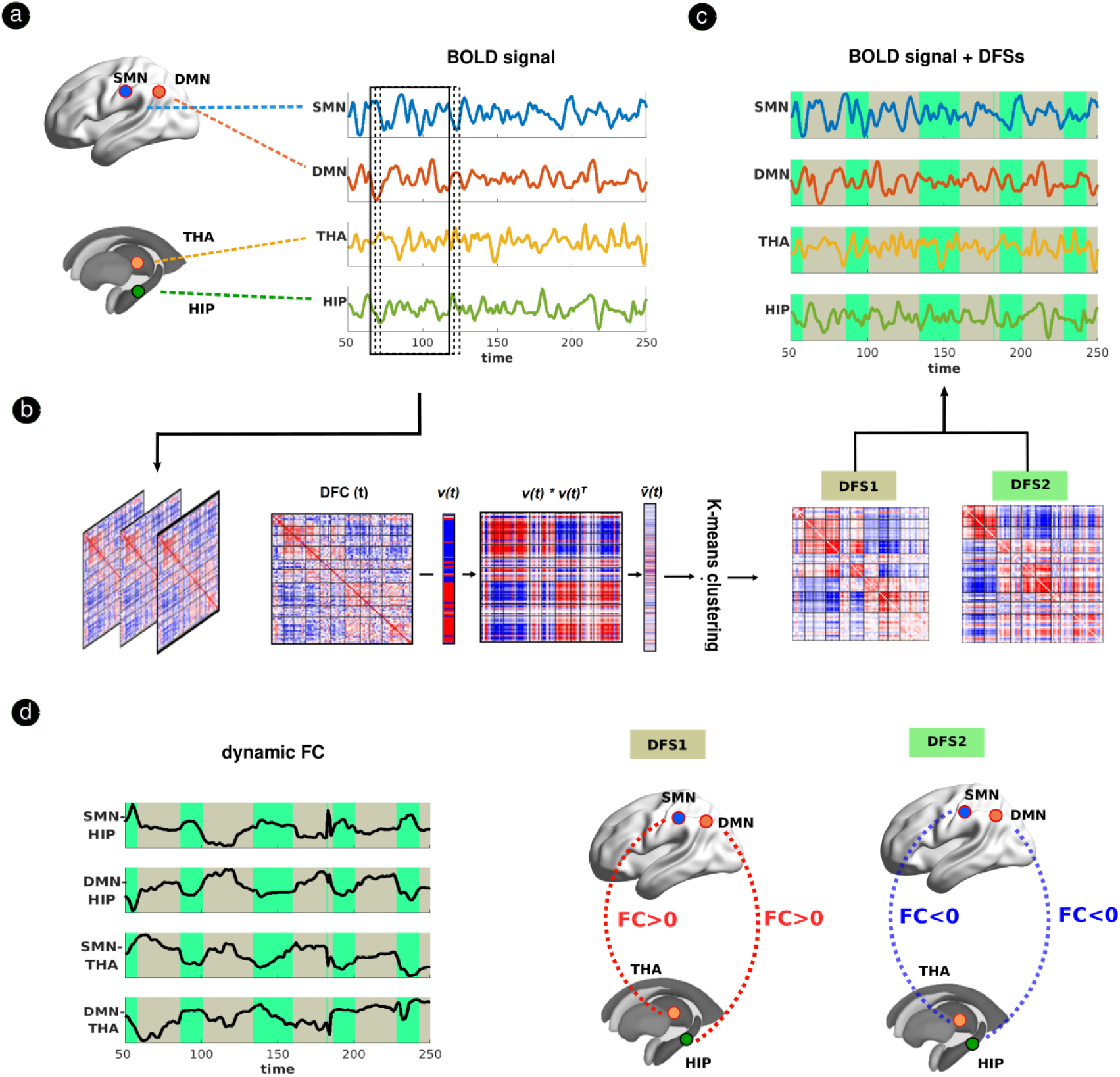
Analysis overview. This work focuses on dynamic cortico-subcortical interactions. As shown by FA22, two groups of subcortical regions (a ‘limbic’ group comprising hippocampus/amygdala and a ‘subcortical nuclei’ group comprising thalamus/basal ganglia) couple dynamically with cortical regions, showing flexible connectivity with task-positive regions (sensorimotor/dorsal attention network) and task-negative regions (default mode network). Connectivity switches are well captured by ‘dynamic functional states’ (DFSs), i.e., recurring patterns of whole brain (cortical-subcortical) connectivity. **(a)** (Average) BOLD signal from four regions belonging respectively to the sensorimotor network, the default mode network, the limbic group and the subcortical nuclei group for an example subject. **(b)** Overview of the analysis pipeline. Sliding window functional connectivity (swFC) is computed using sliding windows of 60 s duration (with a step of 3 s). Then, each swFC matrix is approximated as 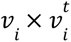, by projecting on the leading eigenspace defined by the first eigenvector v_i_. The upper triangular part of these swFC matrices is vectorized and concatenated across windows and subjects, in order to finally apply a timewise K-means clustering algorithm with correlation distance to identify a set of recurring swFC patterns or DFSs. Each sliding window is assigned to a specific DFS. **(c)** BOLD signal of four key regions (same as in panel a), with different colors highlighting the DFS of the corresponding window (the window centered at that point). Two DFSs capture the dynamic coupling between subcortical and cortical regions. In particular, in DFS1 the hippocampus couples positively with the default mode network and negatively with the sensorimotor network, while the thalamus shows an opposite trend. In DFS2, this pattern of subcortical-cortical connectivity is reversed **(d)**. Time courses of the subcortical-cortical swFC, shaded with different colors according to the corresponding DFSs. The dynamic coupling described above can be noted: switching trends of subcortical-cortical connectivity are summarized graphically in the brain plots on the right.

### Existence of two groups of subcortical regions

FA22 identified two groups of subcortical regions exhibiting anticorrelated FC fluctuations: a first group (‘subcortical group 1’, ‘SC1’) comprising hippocampus and amygdala, and a second group (‘subcortical group 2’, ‘SC2’) comprising thalamus, basal ganglia and cerebellum. This result was obtained by performing a principal component analysis (PCA) on the subcortical projection of *ν* (the first principal eigenvector of the windowed FC, of which we consider entries corresponding to subcortical regions). The rationale behind this analysis was that *ν* can be thought of as a ‘condensed’ representation of the windowed FC.

Consequently, performing a PCA on the subcortical projection of *ν* allows identifying subcortical regions having similar *fluctuations* in windowed FC across time (windows). Two main PCs were found, projecting respectively on SC1 and SC2.

We replicated this analysis in the HCP data set, identifying a main principal component (PC1), explaining 34% of the total variance (Fig. 2a) [2nd component: 14%; 3rd component: 7%; all other components < 4%]. PC1 loaded positively on cerebellum, thalamus, putamen, caudate (bilaterally) and the brain stem; it loaded negatively on hippocampus, amygdala, nucleus accumbens (bilaterally) and had weak loadings on globus pallidus and diencephalon (bilaterally). PC1 is also displayed in a volumetric brain representation in Fig. 2b. This principal component aligned with the subcortical cluster division identified by FA22, as it loaded mostly positively on regions identified as SC1 in FA22, and mostly negatively on the majority of regions identified as SC2 in FA22. Using PC1 loadings, we can thus obtain a division of the subcortex and cerebellum into two groups, which we also term SC1 and SC2. Even though the main split thalamus/basal ganglia vs limbic regions (hippocampus/amygdala) is strongly confirmed by our replication study, we report two discrepancies: globus pallidus was included in SC1 in FA22, but it could not be included in either SC1 or SC2 in our study (it exhibits a weak loading on PC1); the nucleus accumbens was not included in either SC1 or SC2 in FA22, while it was included is SC2 in our study (loading negatively on PC1).

**Fig. 2.**
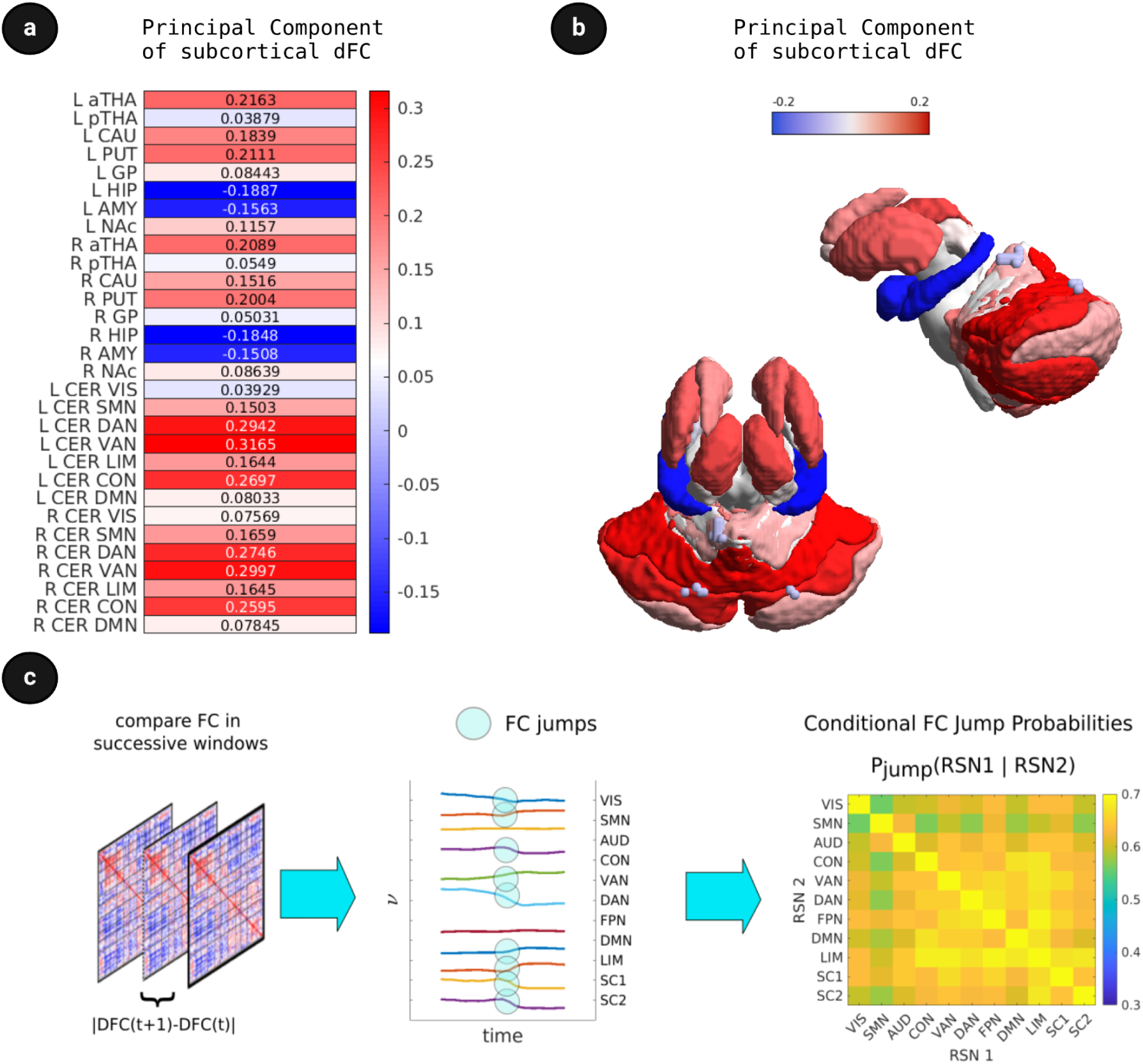
Existence of two main clusters of subcortical regions. We applied PCA on the time evolution of *ν* (principal eigenvector of the sliding-windows FC), restricting attention to subcortical regions **(a)** The first PC shows the competitive relationship between two different subcortical clusters: SC1, which includes the basal ganglia, thalamus and cerebellum, and SC2, which includes the limbic nuclei. CER: cerebellum; THA: thalamus; CAU: caudate nucleus; PUT: putamen; GP: globus pallidus; BST: brain stem; HIP: hippocampus; AMY: amygdala; NAc: nucleus accumbens; DIE: diencephalon **(b)** Visualization of the first PC using a volumetric representation of the brain **(c)**. We computed a metric of “FC change” within each resting state network (RSN) by computing the change of *ν* (principal eigenvector of the sliding-windows FC) across adjacent time windows (left). For each RSN, we record an “FC jump” whenever the change of *ν* is in its 5th upper percentile (middle; blue circles). We then compute conditional probabilities of observing jumps in each RSN, conditioned on concurrent jumps in other RNSs (right).

To explore whether a more fine-grained parcellation would give a more nuanced picture (particularly for the thalamus and cerebellum), we repeated the analysis with different subcortical parcellations. In Fig. S1 we report the results obtained using two subcortical parcellations^20^, one with 8 bilateral regions (‘TianS1’) and the other with 27 bilateral regions (‘Tian S4’). The Tian S1 parcellation roughly aligns with the Freesurfer parcellation, but it excludes some regions (cerebellum, brainstem, and diencephalon) and it splits the thalamus in an anterior and posterior portion. Using this parcellation, we found a main principal component explaining 33% of the variance [2nd component: 15%; 3rd component: 8%; 4th component: 6%, all other components < 5%]. Results are consistent with those obtained with the Freesurfer parcellation, with two main differences. Firstly, the nucleus accumbens is no longer associated with SC2, as it exhibits weak values of PC1. This outcome is more consistent with FA22. Moreover, we a neat split of the thalamus is observed: the anterior portion is strongly associated with SC1, while the posterior one has weak values of PC1, and it cannot be clearly included in either SC1 or SC2. Results obtained with the Tian S4 parcellation (which provides further subdivisions of Tian S1 regions) were very consistent.

Specifically, we found a main PC (PC1) explaining 24% of the total variance [2nd component: 8%, 3rd component: 5%, all other components < 4%]. PC1 loaded positively on regions of the anterior thalamus, caudate and putamen; negatively on regions of the hippocampus and amygdala; and weakly on regions of the nucleus accumbens, posterior thalamus and globus pallidus. To further confirm the split between the anterior and posterior portion of the thalamus, we additionally considered the anatomical Morel atlas of the thalamus^21^, which provides divisions of the thalamus into nuclei (at the coarsest granularity level, the thalamus is split into 7 bilateral portions). We combined the Tian S1 parcellation with the Morel atlas (replacing the anterior/posterior thalamus in Tian S1 with the 7 regions of the Morel atlas, and maintaining the other subcortical regions as given in Tian S1). Results are consistent. We found a main principal component explaining 28% of the total variance [2nd component: 10%, 3rd component: 6%, all other components < 4%]. PC1 loads positively on the medial, lateral and anterior regions of the thalamus, and more weakly on the posterior regions and the additional nuclei (Fig. S1). Finally, we considered a possible subdivision of the cerebellum into 7 regions^22^ associated (by strong values of static FC) with one of the classical seven cortical RSNs^23^. Adding cerebellar regions to the Tian S1 regions, we obtained results consistent with the previous ones. PC1 loads positively on anterior regions of the thalamus, and more weakly on the posterior regions and the additional nuclei (Fig. S1). In summary, this component is very robust and found independently of the specific subcortical parcellation used.

### Coordination of cortical and subcortical connectivity shifts

Another key finding in FA22 was the general coordination between cortical and subcortical connectivity shifts. Mirroring the analysis in FA22, we computed the average windowed FC within each RSN (approximating the FC with its first eigenvector and taking the average of the absolute values within each RSN), and computed an RSN-wide FC change as the difference between two successive windows. We identified an ‘FC jump’ whenever this difference falls in the upper tail of the corresponding distribution (5th percentile). Finally, we computed conditional probabilities *P*(*RSN*;1|*RSN*2) of observing an FC jump in RSN1, given that a jump is observed in RSN2. Synchronization between cortical and subcortical jumps implies a higher-than-chance conditional probabilities of cortical given subcortical jumps and vice versa. All probabilities were >50% (Fig. 2c), implying that FC jumps are strongly synchronized among all regions. Notably, conditional probabilities involving cortical networks and subcortical groups being all >60%. In particular, among the strongest conditional probabilities are those involving simultaneous jumps between SC1 (basal ganglia/thalamus) and cortical networks.

### Dynamic functional states

To find dynamic functional states (DFSs), we performed K-means clustering for increasing values of K (the total number of clusters), using the GordonLaumann+FreeSurfer parcellation^17,18,19^. As typical for K-means, increasing K leads to the survival of the most robust clusters and the splitting of the dimmer into subclusters. The ‘optimal’ number of clusters (assessed though the Silhouette coefficient) should correspond to the largest silhouette value (Fig. S5). Unsurprisingly, the two states found at K=2 (best choice of K) are also the two most stable states across the choice of K. In Fig. 3a/b, we show these two states displayed in a matrix and in a brain surface/volume representation respectively. DFS1 displays high DAN/DMN segregation, positive limbic-DMN connectivity and negative limbic-DAN connectivity, closely resembling the typical pattern of healthy static FC. DFS2 presents a significant DAN/DMN integration, negative limbic/DMN connectivity and a negative coupling between cognitive clusters and sensorimotor clusters. These two states capture the competitive relationship between basal ganglia/thalamus (SC1) and limbic nuclei (SC2), with DFS1 showing a positive correlation between DMN and SC2 and DFS2 showing the opposite FC pattern. Additionally, DFS1 maintains a FC profile similar to healthy static FC. The correlation between cortical and subcortical regions in the two states is summarized in Fig. 3c.

**Fig. 3.**
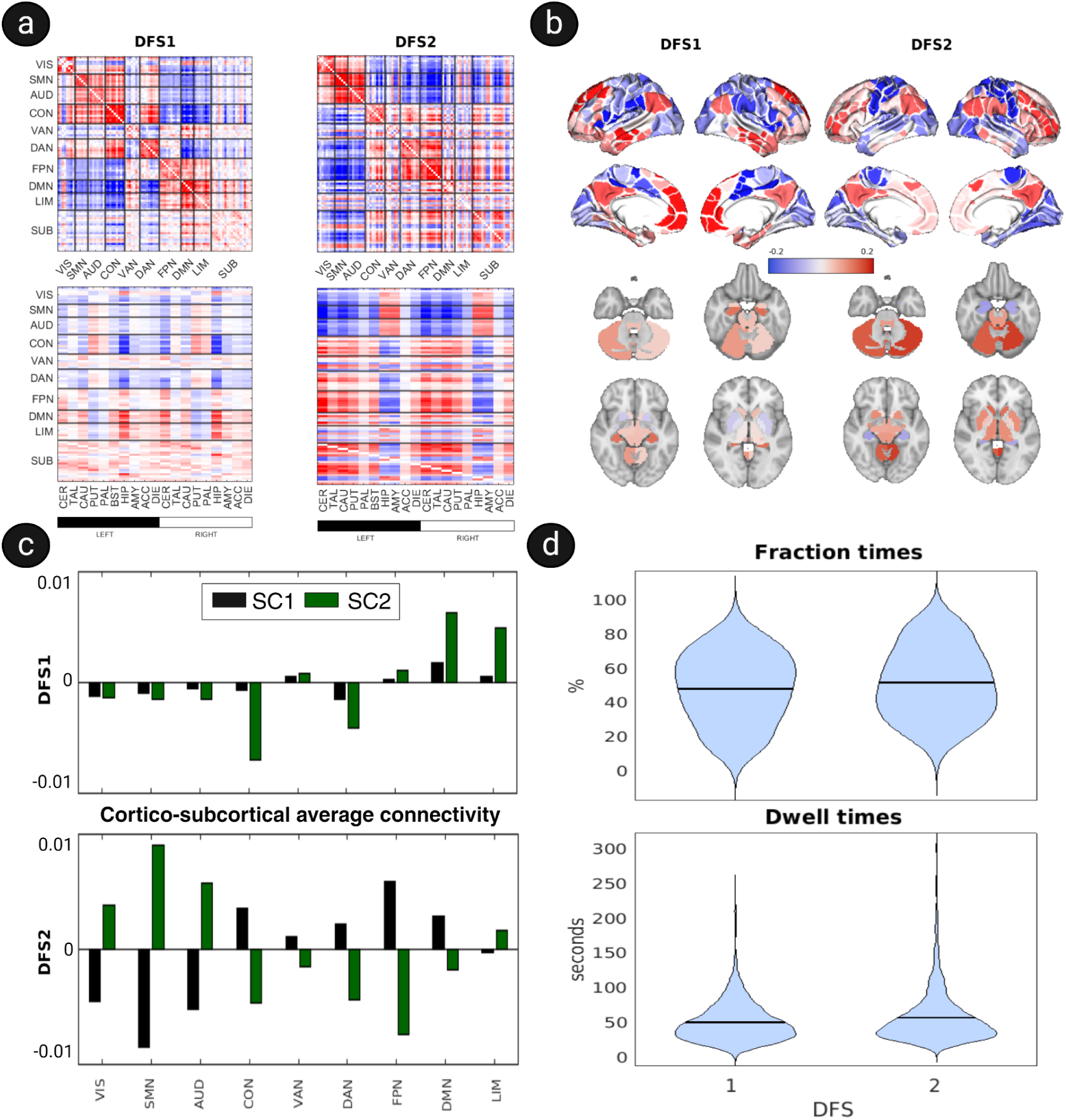
DFS analysis. **(a)** Matrix representation of the Dynamic Functional States displayed globally (top) and with a focus on the cortico-subcortical interactions (bottom). The most robust states observed for K=2 capture the alternating connectivity pattern observed in the Washington dataset between limbic regions (i.e., hippocampus and amygdala) and task-negative (default mode) vs. task positive (dorsal attention and sensorimotor) networks. **(b)** Volumetric representation of the Dynamic Functional States (K=2) **(c)** Average connectivity between subcortical clusters (SC1 and SC2) and cortical networks in the two most robust DFSs **(d)** Distribution of fraction and dwell times for the K=2 states. FTs/DTs was averaged within subjects and then plotted across subjects, resulting in violin plots that display the mean values (black lines). VIS: visual network; SMN: sensorimotor network; AUD: auditory network; CON: cingulo-opercular network; VAN: ventral attention network; DAN: dorsal attention network; FPN: frontoparietal network; DMN: default mode network.

We compared results for K=5 with the 5 states found by FA22 in the Washington University data set (‘WU’). We only partially replicate the patterns found in the previous study. Fig. S6b shows a confusion matrix representing the similarity of the K=5 states found by FA22 and the states found in the HCP data set (‘HCP’). Similarity is assessed by the correlation between the state centroids. The DSF1 (WU) is approximately reproduced by the DFS1 (HCP), with a correlation of 0.82. This state corresponds to the DFS1 found with K=2. The DFS3 (WU) is well reproduced by the DFS4 (HCP; correlation value of 0.7), and partially reproduced by the DFS1 and DFS3 (HCP; with correlation values of 0.5 and 0.46 respectively). DFS4 (WU) resembles (the correlation value is 0.68) the DFS3 (HCP), as they both show a weak cortico-subcortical connectivity. DSF2 and DFS5 (WU) could not be associated with any of the DFSs found in HCP. Notably, DFS2 (WU) was particularly common in stroke patients at the acute stage, showing abnormal integration of cortical connectivity. Comparable fraction times and dwell times are observed for both data sets.

We tested the robustness of the K=2 analysis with respect to preprocessing and parcellation steps. We replicated the analysis by changing the cortical and subcortical parcellation used. For a different cortical parcellation, we used the well-known Schaefer atlas with 100 regions^24^. For the Schaefer atlas, regions are divided into RSNs according to the classical classification^23^, where the control network (CON) includes regions associated to the frontoparietal network and the VAN includes regions associated to the cingulo-opercular network^17^. For a different subcortical parcellation, we used the same atlas described above^20^, which does not include cerebellar regions. In Fig. S7 we show the results of changing cortical and subcortical parcellation, respectively. The core structure of the DFS (K=2) is preserved across the parcellation changes. In particular, DFS1 is characterized by a strong DAN-DMN anticorrelation, while the hippocampus couples positively with the DMN and negatively with the task positive networks (DAN and primary networks). Conversely, DFS2 is characterized by a strong segregation of the primary networks from the association networks. Hippocampus/amygdala (SC2) correlate positively with primary networks and negatively with association networks, while the opposite pattern is observed for thalamus/basal ganglia (SC1). Next, we investigated the impact of specific preprocessing steps. We first addressed the impact of the temporal downsampling step (used to match the TR of previous studies, TR=2). Downsampling had virtually no effect on the clustering results (Fig. S3). We then tested the impact of global signal regression (GSR). Omitting GSR produced a substantial change in the structure of DFSs (Fig. S2), confirming the sharp effect of this preprocessing step. Finally, to test whether the lagged cross-covariance structure of the data is sufficient to yield the observed DFSs, we applied phase randomization (PR). PR generates surrogate data that preserve the empirically measured cross-covariances but are linear and Gaussian^25^. As shown in Fig. S4, DFSs resulting from the PR-generated time series closely resemble the ones obtained with the original data. This correspondence is furtherly investigated in the confusion matrix represented in Fig. S4b. The latter shows a neat one-to-one correspondence between the two sets of clusters, with very similar centroids.

### Correlation with behavior

We tested a possible correlation between individual dynamic FC metrics and individual behavioral traits. The large array of behavioral variables (“subject measures”) available in the HCP data set were summarized into a few general descriptors capturing key aspects of cognition and behavior (Fig. 4c). We first used the positive-negative mode (PNM) of behavior-FC covariation^26^, which is a single indicator of global behavioral/social function (Methods). In addition, we considered seven individual markers identified in a recent study^27^ which capture different aspects of cognition (Fig. 4c).

**Fig 4.**
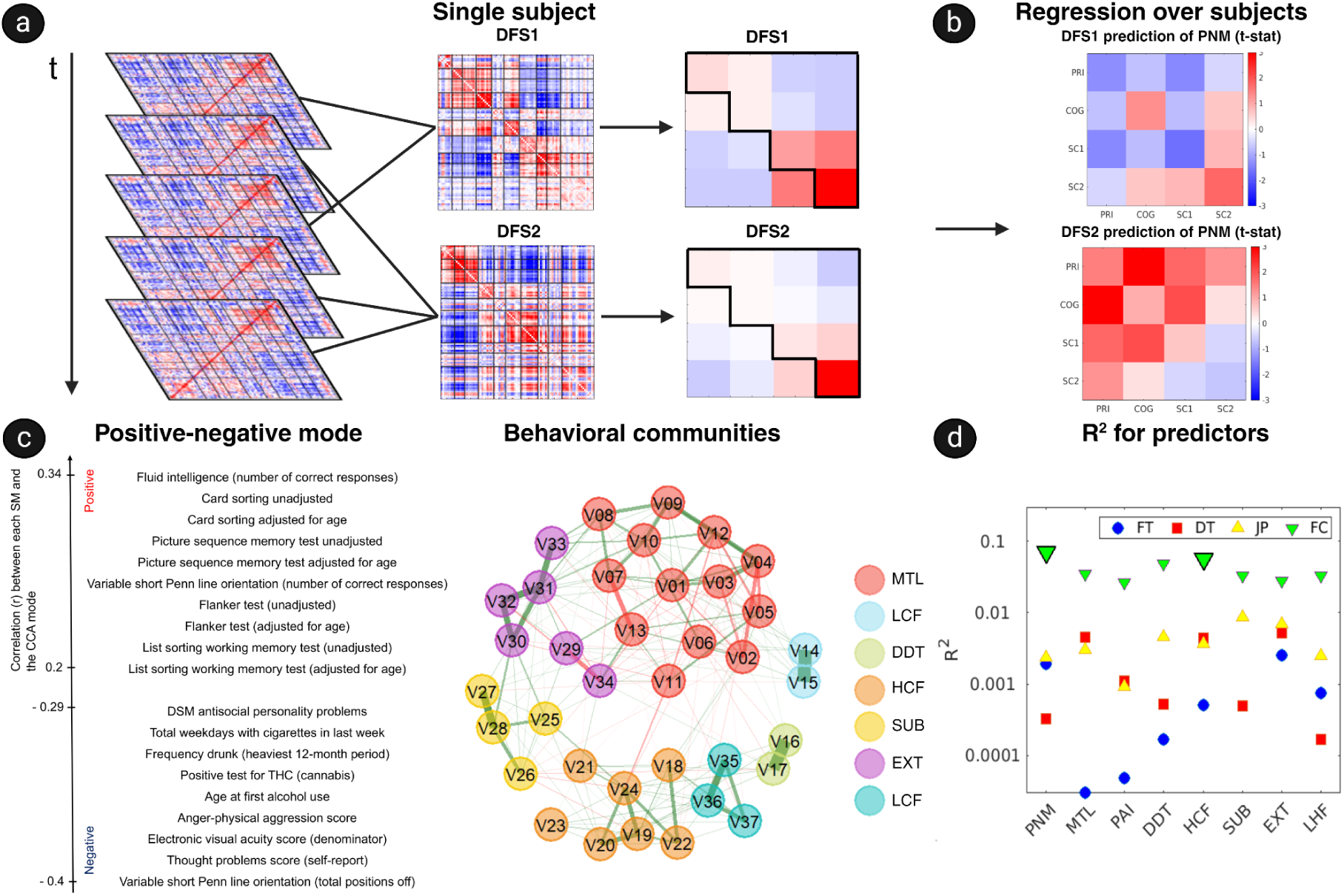
Functional connectivity and behavior. We investigated the impact of functional connectivity differences among individuals on behavior by carefully scrutinizing the relationships between different facets of the former and the latter through a series of Generalized Linear Models (GLMs). **(a)** For each subject we separately averaged the windows assigned to DFS1 and DFS2 by the clustering procedure, resulting in an individual representation of the states. These representations were reduced in dimensionality to 4x4 matrices (of which we retained the upper-triangular part) averaging entries pertaining to primary networks (PRI), cognitive networks (COG) and the two subcortical components SC1/2. **(b)** We show t-statistics associated with the coefficients of the linear model trying to predict the PNM from individual 4 x 4 connectivity patterns in the two DFSs. **(c)** In order to encompass the complexity of the multitude of behavioral and demographic indices available thanks to the HCP consortium, we employed two alternative methodologies: the first one^26^ established a behavioral positive-negative mode (PNM) allowing to determine a score for each participant depending on their overall performance in a series of 150 behavioral/demographic tests (for a complete list see Supplementary Information). The second one^27^ grouped 38 behavioral/demographic indices into 7 clusters: mental health (MTL), pain (PAI), low cognitive functions (LCF), delay discounting (DDT), high cognitive functions (HCF), substance abuse (SUB), externalizing problems (EXT). From each community we extracted a score for each subject. **(d)** We then considered several aspects of FC as predictors (fraction/dwell times, P(jump in area1|jump in area2), individual DFSs) in an exploratory analysis to determine the most relevant representations by the means of their coefficient of determination (magnified and highlighted in black).

We first tested for correlation between behavioral metrics and individual fraction/dwell times obtained with K=2. We performed a linear regression using the behavioral metrics as dependent variables and the fraction/dwell times as predictors. The total regression R^2^ was always lower than 0.01, meaning that fraction/dwell times explain less than 1% of the variation in the behavioral metrics considered (Fig. 4d). We thus found no significant effect of fraction/dwell times on behavioral metrics (permutation test on R^2^, corrected for 32 comparisons). In addition, we tested whether the probabilities of synchronized jumps between cortical regions and the two main subcortical clusters could predict behavior. We averaged the conditional probabilities (Fig. 2c) by aggregating cortical regions, regions belonging to SC1, and regions belonging to SC2, obtaining a 3 x 3 matrix of synchronized jumps. From this matrix we extracted four entries corresponding to the probabilities of synchronized jumps between the cortex and, respectively, SC1 and SC2. We performed a linear regression using the behavioral metrics as dependent variables and these probabilities as predictors. The total regression R^2^ was always lower than 0.01 and non significant (Fig. 4d). Finally, we tested whether the average FC patterns in each DFS could predict behavior. We computed an average DFS pattern for each individual subject, by averaging the sw-FC over time windows assigned to one of the K=2 DFSs (Fig. 4a). This can be considered as a DFS pattern ‘adjusted’ to each participant. We averaged the FC patterns by aggregating task-positive regions (SMN, DAN, VIS), task-negative regions (CON, DMN), regions belonging to SC1, and regions belonging to SC2, obtaining two 4 x 4 matrices (one for each DFS) for each participant. We performed a linear regression using the behavioral metrics as dependent variables and the entries of this matrix as predictors. The total regression R^2^ was significant for the PNM, which displayed R^2^ = 0.07, with a significant effect (P=0.003, permutation test on R^2^, corrected for 32 comparisons). Higher values of the PNM are associated with higher segregation within the cortex, and between the cortex and SC1 in DFS1; and, conversely, higher integration within the cortex, and between the cortex and SC1 in DFS2 (Fig. 4b).

## DISCUSSION (1954)

Human neuroscience has traditionally focused on the neocortex, often neglecting the subcortical brain. This ‘corticocentric bias’^28^ reflects both technical limitations in imaging the subcortex and the widely held misconception that ‘higher’ cognitive functions in humans would mostly depend on the neocortex, considered as the culmination of brain evolution. Research in the last decade has shown that ‘ancient’ subcortical regions underwent significant reorganization accompanying cortical expansion during evolution^29^, and they play a critical role in advanced cognitive functions^30,31,32^. A recent proposal^5^ holds that cognition would rest on a highly parallel, ‘shallow’ neural network architecture whereby hierarchical cortical processing is complemented by cortical-subcortical loops. While these considerations would suggest systematically including subcortical regions in the description of large-scale brain dynamics, only few studies have focused on whole-brain cortico-subcortical functional connectivity (FC) so far. The majority of these works portray a *static* coupling between cortex and subcortex, trying to ‘affiliate’ subcortical structures to well-known cortical resting state networks through their static FC^33,34,35,36,37^. An exception is FA22, where *dynamic* functional connectivity was investigated in a mixed sample including mainly stroke patients, providing evidence that subcortical structures couple flexibly with cortical networks. In the present work, we focus on a much larger sample of healthy young subjects, the Human Connectome Project HCP1200 data set. Our results provide substantial support for the main conclusions of FA22.

In FA22, the dynamic FC analysis identified two main clusters of subcortical regions. The first (‘subcortical cluster 1’ or ‘SC1’) comprised the thalamus, basal ganglia and cerebellum; the second (‘subcortical cluster 2’ or ‘SC2’) the hippocampus and the amygdala. These two clusters are already distinguishable at the level of subcortical static FC (Fig. S8). Regions within the two clusters show positive FC, while negative FC is observed between the two clusters. Fine-grained parcellations of the thalamus and cerebellum confirm this ‘two-block’ picture with minor refinements (the posterior thalamus and the ‘visual’ cerebellum show only weak affiliation with SC1). In terms of (static) cortical FC, SC1 shows weakly negative coupling with primary regions (SMN, AUD, VIS) and weakly positive coupling with associative networks (CON, DAN, VAN, FPN, DMN). On the other hand, SC2 shows a strongly positive FC with task-negative regions (DMN), a weakly positive FC with primary networks, and a weakly negative FC with associative networks. However, our dynamic FC analysis reveals that static FC ‘hides’ a rich dynamic picture where the two subcortical clusters couple flexibly with cortical regions. In particular, the two clusters also have anticorrelated *fluctuations* in FC. When doing a principal component analysis on sliding-window FC, we identified a first principal component with positive loadings on SC1 and negative loadings on SC2 (Fig. 2a), consistently across different subcortical parcellations (Fig. S1). The two clusters thus seem to behave as ‘cohesive blocks’ in terms of their static and dynamic FC. The internal cohesion of these two clusters is in agreement with both classical and recent results. SC2 comprises regions conventionally considered part of a single system, the ‘limbic system’ for emotion and memory^38,39^. The tight coupling within

SC1 is less straightforward to interpret. The cerebellum and basal ganglia were traditionally thought to be independent, giving complementary contributions to learning and motor control^40^, and to communicate only at the cortical level. However, recent findings provided solid evidence that the two systems are reciprocally interconnected not only at the level of the thalamus^41^, but also through more direct subcortical pathways^42,43^. This suggests that cerebellum, basal ganglia and thalamus constitute an integrated network^42^ that acts in concert with the cortex via cortical-subcortical loops. On the contrary, the apparent antagonism between the two clusters is neither widely discussed nor reported in the literature. An exception is a recent contribution^43^, where cortical-subcortical interactions were investigated with edge-centric FC, showing that edges involving hippocampus and amygdala have a distinct pattern from those involving striatum and thalamus.

FA22 showed that large shifts in the connectivity of cortical regions are accompanied by large shifts in the connectivity of subcortical ones, providing evidence that FC rearrangements are a phenomenon involving the cortex and the subcortex jointly. Here we confirm this picture, observing that subcortical connectivity shifts predict cortical shifts with >60% accuracy and vice versa (Fig 2c). One may speculate that FC shifts are *driven* by subcortical regions, but possible mechanisms remain very hypothetical. Invasive recordings in rats have revealed that different types of subcortical activity, such as slow-frequency activity^45^, ripples^46^ and dentate spikes in the hippocampus^47^, regular spikes^48^ and spindles^49^ in the thalamus can trigger widespread cortical effects. In two cases, simultaneous electrophysiological/fMRI recording allowed observing subcortical effects on the cortical BOLD signal^45,49^, such as FC changes triggered by hippocampal events^45^. However, no such evidence has been collected in humans so far. Given the difficulty of performing causal manipulations and invasive recordings in humans, future work could at least characterize directional interactions between subcortical and cortical areas^50^ through Granger causality or effective connectivity^51^.

Simultaneous shifts in cortical-subcortical FC arrangements are well captured by dynamic functional states (DFSs). Our analysis identified K=2 as the optimal number of states, and, congruently, identified two states that are consistently observed across different values of K (Fig. S5). The first state (DFS1) is characterized by a strong DAN-DMN anticorrelation, and a correspondingly antinomic pattern in the coupling of SC2 (hippocampus/amygdala) with the cortex: SC2 couples positively with the DMN and negatively with the task positive networks (DAN and primary networks). Conversely, the second state (DFS2) is characterized by a strong segregation of the primary networks from the association networks. SC2 correlates positively with primary networks and negatively with association networks, while the opposite pattern is observed for SC1. Thus, these two states capture the competitive relationship between SC1 and SC2, and their flexible coupling with task-negative vs. task-positive regions.

Currently, we hesitate to advance strong hypotheses on what could drive these state alternations. However, recent literature suggests arousal as one possible mechanism, given its synchrony with FC changes^11^. Based on concurrent fMRI-pupillometry or fMRI-EEG, clear evidence was given that arousal can modulate FC ^52,53,54,55,56,57,58^. In particular, some works related arousal variations with FC dynamics, identifying ‘low-arousal’ and ‘high-arousal’ states. Decreases in arousal were associated with decreases in anticorrelation between the default mode and task-positive networks^52^, and a low-arousal state characterized by negative FC between the thalamus and primary networks and reduced DAN-DMN anticorrelation was identified^55^. This pattern shows remarkable similarities with DFS2 in the present study. These similarities suggest that arousal could contribute to DFS switches we observe. If arousal plays a role, then it would not be surprising to observe an involvement of subcortical regions in state switches, as the role of the thalamus in the regulation of arousal is well recognized^59^. In this context, our findings would suggest that thalamus-mediated changes in arousal have a large impact on the organization or cortical and limbic connectivity.

While we qualitatively reproduced the main results of FA22, there are some discrepancies between the set of DFSs found in FA22 and those found in the present work. Using a very large sample (n=7500), Abrol et al.^60^ reported typical similarities above 0.8 between the cluster centroids of DFSs obtained in different subsamples - a stronger consistency than the one observed between the present study and FA22. There are several possible explanations for this difference. First, the cohort studied in FA22 mostly included stroke patients. The latter generally present widespread alterations of FC^61^, such as reduced interhemispheric FC and increased intra-hemispheric FC. One of the DFSs found in FA22 (DFS2) corresponded to this stereotypical ‘stroke’ pattern, and it is therefore unsurprising that we miss it in the present study. Other differences (such as the absence of a pattern corresponding to the DFS5 in FA22) may be due to additional discrepancies between the data sets. The two cohorts markedly differed in terms of average age (53 years for FA22, 23 years for HCP), a variable that has a large influence on the functional connectome^62^. Acquisition was not identical (TR=0.7s in this study, TR=2s in FA22), and preprocessing pipelines exhibit minor differences (nuisance regression was performed through ICA-fix for HCP and regression of nuisance time series in FA22).

Testing for “significance” of the observed DFSs is very hard, as there is no consensus on the appropriate null model to use in this case. While phase randomization (PR) has been often employed to test for dynamic connectivity^25^, PR surrogates match all spectral and cross-correlation properties as the original data (power spectra and cross-covariances at all lags are preserved), and it is quite contentious that they can be genuinely considered to lack FC dynamics^63^. Many dynamical features observed in fMRI times, such as a large variability in sw-FC^64^, co-activation patterns^65^, or ‘events’ of high-amplitude co-fluctuation^66^, also appear in PR surrogates^25,67,68^. In fact, while PR data are stationary (in the statistical sense), this does not imply that observed window-to-window FC variations are just sampling noise, but merely that such variations do not follow a well-defined temporal trend. Our observations of closely matching DFSs in PR surrogates is in line with previous literature^60,63^, with which we share the main conclusion, i.e., that the emergence of DFSs substantially depends on the lagged cross-covariance structure of the time series.

We investigated whether dynamic FC can predict aspects of individual cognition or behavior. Looking for statistically reliable association between a large set of behavioral variables (subject measures, SMs) and brain function may require very large samples^69^. Therefore, despite the availability of 262 variables across 15 behavioral domains in the HCP data set, we restricted attention to a few summary metrics effectively summarizing several SMs. The first was the ‘positive-negative mode (PNM) of population covariation’^26^, highlighting a global individual ‘function outcome’ associated with cognitive function, emotion regulation, alcohol and substance use. We then considered seven ‘factors’^27^ summarizing different cognitive domains. We did not find a significant relationship between SMs and DFS metrics such as fraction times (FT) and dwell times (DT), with all R^2^ < 0.005. Analogously, we did not find a relation between behavior and the probabilities of cortical/subcortical FC jumps (R^2^ < 0.01). Consistently with our findings, Lee et al.^70^ previously reported weak correlations of behavior with DT/FT co-activation patterns. These results suggest that temporal summary metrics of dynamic FC, such as FT/DT and jump probabilities, do not significantly contribute to explaining inter-individual behavioral variability. This is possibly due to the fact that such summary measures do not account for variation in the strength of specific FC links. A link-wise analysis of static FC^26^ explained a large fraction of the PNM variance (R^2^=0.75). Similarly, a significant fraction of behavioral variability in the HCP data could be predicted using all link strengths in the non-lagged and 1-lagged FC matrices^71^. When considering network-averaged FC strengths associated with DFS1 and DFS2 in single individuals, we obtained significant correlations, up to R^2^=0.07 for the PNM and a measure of. In particular, more positive values of the PNM (generally associated with better cognitive health) corresponded to a dynamic cortical-subcortical connectivity pattern, where cortex and SC1 are *less integrated* in DFS1, and *more integrated* in DFS2. Healthier cognition may thus require an alternation of states with higher and lower cortico-subcortical integration.

In conclusion, our study replicates the main findings of FA22. The human brain at rest is characterized by large fluctuations in functional connectivity, with synchronized changes occurring in the cortex and the subcortex. Connectivity oscillates between states corresponding to different patterns of cortical-subcortical connectivity, captured by different dynamic functional states (DFSs). Two main groups of subcortical regions, one comprising the thalamus, basal ganglia and cerebellum, the other comprising limbic regions such as hippocampus and amygdala, show flexible coupling arrangements with task-positive and task-negative cortical regions. The mechanisms underlying synchronous cortical-subcortical connectivity changes are presently unknown and demand further investigation, possibly integrating neuroimaging results with electrophysiologic and behavioral measurements.

## MATERIALS AND METHODS

### Washington dataset

All subjects included in FA22 were scanned with a 3T Siemens Tim-Trio scanner at the Washington University School of Medicine with a standard 12-channels head coil. They were divided into four experimental groups: three groups of stroke patients, depending on the time they were scanned after stroke onset (1-2 weeks, 3 and 12 months after) and one age-matched control group. Pulse sequence included a gradient-echo EPI sequence with TR=2s acquiring 32 contiguous 4 mm slices, with 4x4 mm in-plane resolution while fixating on a small white crosshair. Pre-processing included: regression of head motion, signal from ventricles and Cerebrospinal fluid, signal from white matter, global signal; temporal filtering retaining frequencies between 0.009 and 0.08 Hz; frame censoring, with framewise displacement of 0.5 mm. After all the pre-processing steps, a total of 20 controls and 47 patients with first-time strokes were considered for the analysis.

### Human Connectome Project dataset

The HCP’s dataset included 1206 participants, who underwent neuroimaging sessions and a large battery of behavioral tests. Of the 1206 participants, 1096 were scanned with a modified 3T Siemens “Connectome skyra” scanner at the Washington University, using a standard 32-channels Siemens receive head coil and a specifically designed “body” transmission coil. Pulse sequence included slice-accelerated multiband acquisition with a multiband factor of 8, spatial resolution of 2 mm isotropic voxels and TR=0.7s. Participants underwent two 15-minutes scanning sessions with opposite phase encoding directions (L/R and R/L), while fixating on a crosshair. We included in the analysis only participants that were scanned both in the L/R and in the R/L direction for 840 s (n=1078). We used pre-processed data provided by the HCP. The HCP’s preprocessing pipeline is divided into two distinct protocols^72^: one applied entirely on the volume data involving temporal filtering and de-noising and the second one regarding mapping the data to cortical surfaces and subcortical gray-matter domains using the Connectivity Informatics Technology Initiative file format (CIFTI). One promising approach for removing structured artifacts involves denoising each 15-minutes rfMRI scan with the Independent Component Analysis (ICA) based tool called FSL’s MELODIC. This tool, paired with the FMRIB’S ICA-based X-noise filter, allows decomposing the data into multiple components (comprising a spatial map and a corresponding time course) and to classify them in order to subsequently regress out the confounding ones. Additionally, in line with FA22, we included two supplementary pre-processing steps: signals were band-passed in the frequency band [0.009 Hz,0.08 Hz] with a Butterworth filter of order 1 and the mean GM signal was linearly regressed (global signal regression).

### Parcellations

For our initial analysis, we used the same parcellation used in FA22. Time series were projected on the cortical surface of each subject divided according to the resting state functional connectivity boundary mapping developed by Gordon et al.^17^. This technique leverages abrupt transitions in resting-state functional connectivity (RSFC) to noninvasively identify the borders separating cortical areas. The original parcellation includes 333 regions, but all regions with <20 vertices (∼50 mm2) were excluded due to low signal-to-noise ratio (SNR) (Siegel et al., 2016). The remaining 324 regions were further reduced to 71 by a clustering procedure^14^ and grouped into 8 resting state networks (RSNs): Visual Network (VIS), Sensory Motor Hand-Mouth Network (SMN), Auditory Network (AUD), Control or Cingulo-Opercular Network (CON), Ventral Attention Network (VAN), Dorsal Attention Network (DAN), Fronto Parietal Network (FPN), Default Mode Network (DMN), Limbic Network (LIM). We also considered 19 subcortical and cerebellar regions derived from the FreeSurfer subcortical atlas^18,19^. Expanding the initial analysis to aid the investigation of cortico-subcortical interactions, we used a different parcellation of the subcortex^20^, which provides four different parcellations with an increasing degree of granularity. The coarsest parcellation includes 8 bilateral regions, while the most fine grained one comprises 27 bilateral regions (see Supplementary Table 2 of Tian et al.^20^). This subcortical cartography was based on resting state functional connectivity gradients (“gradientography”): region boundaries were identified on the basis of strong shifts in functional connectivity gradients. To analyze the cerebellum, we considered the cerebellar parcellation by Buckner et al.^22^. This parcellation was obtained by considering the resting-state FC between the cerebellum and the cortex. In particular, the cortex was divided into seven RSNs ^23^, and the FC between each voxel in the cerebellum and each cortical RNS was assessed; based on the maximal FC, cerebellum voxels were assigned to one of seven clusters based on their maximum correlation with cortical regions. Finally, we considered an anatomical parcellation of the thalamus (‘Morel atlas’) provided by Krauth et al.^21^. This map was obtained from detailed histological maps of the thalamus.

### Sliding-window functional connectivity

FC dynamics were investigated through sliding window temporal correlation, one of the most straightforward approaches for dynamic FC analysis. Similarly to a moving average function, this technique computes a succession of pairwise Fisher z-transformed Pearson correlation matrices, relative to windows of a given width. These correlation matrices are informative of the time-varying FC between the networks considered in the brain parcellation of choice. Importantly, to compensate for the difference in TR durations between the two datasets (i.e., FA22’s TR=2s and HCP’s TR=0.7s), we down-sampled HCP’s timeseries to one third of the points. Then, from the down-sampled timeseries, we extracted windows lasting approximately one minute (28 TRs), with a sliding step of 3 TRs (approximately 2s). Window-length choice represents a critical point in dynamical functional connectivity analysis^73^. Namely, having windows shorter than the analyzed components’ wavelengths might cause spurious fluctuations in dFC. Similarly, too long windows might prevent legitimate functional fluctuations to be identified. Thus, we selected our sliding window’s width on the basis of previous results, matching FA22’s choice. Additionally, each correlation matrix was approximated by projecting it onto the corresponding eigenspace, defined by the first eigenvector *ν*_i_ . Since eigenvectors are defined less than the sign, we averted this problem by translating each eigenvector into the reconstructed square matrix 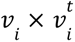, saving the vectorized upper-triangular part alone, avoiding redundancies in the data. Ultimately, all the resulting vectors were concatenated across windows, subjects, and time points.

### Dynamic Functional States’ definition

The last step for defining Dynamic Functional States (DFS), required the application of a time-wise K-means clustering procedure with correlation distance. K-means is a clustering technique, aiming to partition an N-dimensional population into k clusters based on a sample. Each observation belongs to the cluster with the nearest mean (e.g., cluster centroid), serving as a prototype of the cluster. In this case, this procedure resulted in a set of five Dynamic Functional States (all the operations described up to now are summarized in Fig. 1b). This algorithm minimizes within-cluster variances, taking into account a range of possible distance metrics. For this analysis, we employed the correlation distance, which is defined as follows:

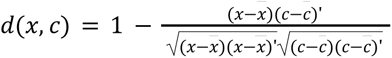

Where: *x* is an observation and *c* is a centroid. In addition, 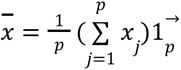, 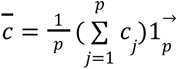 and 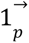 is a row vector of *p* ones. Furthermore, the optimal K value was deducted by comparing the clustering performances with different numbers of clusters (from 2 to 6), with respect to a metric for interpreting and validating the consistency within clusters of data: the Silhouette value. This parameter is a measure of the fitness of a certain data point for its cluster of belonging, compared to other clusters. This metric ranges from -1, indicating the lowest fitness and +1, indicating the highest fitness. Then, for a certain data point *i* ∈ *C_I_*, where *C_I_* is the cluster of belonging, the Silhouette value is defined as follows:

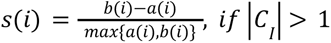

Where, *a*(*i*) is the mean distance between *i* and all the other data points belonging to the same cluster, while *b*(*i*) is the smallest mean distance between *i* and all the data points belonging to other clusters.Additionally, the clustering procedure associated each sliding window to a specific DFS, so that for each subject we had a discrete time series *x*(*n*) (with n ranging from 1 to 746), where each value represented the active Dynamic Functional State for that time window. These time courses allowed us to evaluate three dynamical measures for each state, namely: fraction time *f_k_*, being the percentage of times during which a state is active:

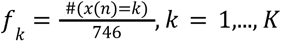

where #(*a*) stands for the number of occurrences of the condition *a* . The dwell time *l_k_*, being the average length of periods in which each state remains continuously active:

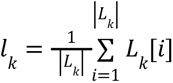

where *L_k_* is the set having cardinality |*L_k_*|, with each element *L_k_*[*i*] representing the length of a period of continuous activity of state *k*. The transition probability *DFS_i_ > DFS_j_*, from *DFS_i_* to *DFS_j_*, where:

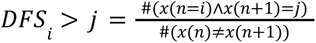

being the ratio between the number of jumps from *DFS_i_* to *DFS_j_* over the total amount of jumps.

### Phase randomization

Phase randomization (PR) is a common framework for generating null data extensively employed in physics^74^. Recently, it has also been applied to fMRI data for studying dynamic FC^75,76^. The PR procedure performs a Discrete Fourier Transform (DFT) of the original time series, adds a uniformly distributed random phase to each frequency, and then performs the inverse DFT to create surrogate data. Crucially, the random phases are created individually for each frequency, but they remain consistent across various regions of the brain. Adding the same random phase to the same frequency components of the RSNs preserves the static FC and the lagged cross-covariance structure in the surrogates (in addition, also the mean, variance and power spectrum of the signals are preserved). This class of surrogates correspond to the null hypothesis that time series are generated by a linear, stationary Gaussian process^25^.

### Behavioral analysis

The HCP provides a large array of subject measures (SMs; i.e. individual measures for each participant), covering demographic, psychometric and behavioral information. The full list of SMs with a detailed description can be found in the HCP 1200 Manual. SMs comprise demographics (e.g., education, employment, income); physical and mental health history, present and past use of tobacco, alcohol, marijuana, and other drugs; Symptoms/history of eating disorders, depression, psychosis, antisocial personality, obsessive-compulsive disorder, post-traumatic stress, social phobia, panic attack; Folstein MiniMental State Exam; Pittsburgh Sleep Quality Index; Parental Psychiatric and neurologic history; Handedness assessment; Menstrual cycle and other endocrine information in females; Urine drug assessment, breathalyzer test, Blood test; NIH Toolbox behavioral tests (which includes 19 sub-domains within the broad domains of cognitive, motor, emotional and sensory functions); Non-NIH Toolbox behavioral tests (color vision, contrast sensitivity, personality, attention, episodic memory, fluid intelligence, emotion processing, spatial processing, and delay discounting)

Starting from a subset of 158 of such SMs, Smith et al.^26^ performed a canonical correlation analysis (CCA) linking the SMs with the individual static FC matrices, which resulted in a principal axis of FC-SM co-variation. They list 59 SMs (listed in Supp. Inf.) with a large loading onto the principal axis. We replicated the analysis by Smith et al.^26^ using the same methodology. However, we used a larger cohort of subjects from the HCP database (i.e. 1206 vs 461 HCP subjects), and we computed connectivity matrices in the Schaefer100 + Freesurfer cortico-subcortical parcellation (while Smith et al.^26^ used a customized functional parcellation with 200 regions). The raw behavioral measures for the selected 158 SMs were initially subject to a rank-based inverse Gaussian transformation to enforce Gaussianity, avoiding the influence of potential outliers. Additionally, 17 potential confound SMs (including head motion) were regressed out from the behavioral data (for a complete list see Smith et al.^26^). To account for missing data, a subjects x subjects covariance matrix was estimated by ignoring missing values for either subject, which was then projected onto the nearest valid positive-definite covariance matrix. Finally, the eigenvalue decomposition was computed onto the resulting covariance matrix, and the first 100 eigenvectors were kept. Regarding connectivity data (referred as *N*), we computed subject-wise partial temporal correlation between the time series of each region keeping only the upper-triangular part of each correlation matrix. The resulting vectors were concatenated across subjects and the Pearson correlation values transformed into z statistics with Fisher’s transformation. Then, this connectivity matrix was demeaned column-wise, globally variance-normalized and the same confounds SMs were regressed out. Lastly, a principal component analysis was computed on *N*, keeping the first 100 components. The fully preprocessed behavioral and connectivity matrices (*S* and *N* respectively) were ultimately fed into a CCA, identifying 100 components aiming to optimize de-mixing matrices *A* and *B* to ensure that the resulting matrices *U* = *N*\* *A* and *V*= *S*\**B* were highly similar to each other.

Granziol and Cona^27^ analyzed 38 SMs reflecting cognitive and processing aspects, mental health and behavioral problems, personality characteristics, and substance use frequencies (the list of the 38 SMs is reported in Supp. Inf.). Exploratory Graph Analysis^77^ (EGA) was used to cluster these SMs into “communities” or clusters of SMs characterized by high correlation. Briefly, EGA works with the following steps: i) the graphical LASSO algorithm^78^ was used to find partial correlations between the 38 SMs ii) the walktrap community detection algorithm^79^ is applied to find clusters/communities. Seven domains were identified (mental health, substance abuse, low cognitive functions, high cognitive functions, pain, delay discounting and externalizing problems).

We replicated the analysis by Granziol and Cona^27^ using the graphical LASSO algorithm with sparsity 0.5, as implemented in the R package ‘glasso’; we then applied the walktrap community detection as implemented in the R ‘igraph’ package. We computed network loadings for each measure as follows^80^: starting from the partial correlation matrix *W_ij_*, where *i*, *j* = 1, …, *N_SM_*, we considered all SMs assigned to factor *c* and computed loading as 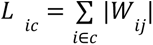 and then we normalized loadings as 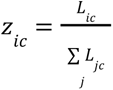. Given the set of SMs for all subjects, *X_ik_* where *k* = 1, …, *N_subjects_*, we computed community/cluster scores for each subject as 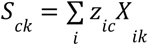.

Ultimately, we assessed the impact of variability in dFC patterns via a Generalized Linear Model (GLM) in an exploratory analysis considering different predictors (fraction times, dwell times, *Pjump*(*RSN*1|*RSN*2) and individual DFSs) and the two behavioral scores described above (i.e. Smith’s behavioral mode and Granziol’s 7 communities scores) as alternative dependent variables.

### Supplementary Data

List of the 59 subject measures (SMs) with significant loading onto the principal axis of Smith et al. (2015), with their unique identifiers as provided by the HCP consortium:

PicVocab_Unadj, PicVocab_AgeAdj, PMAT24_A_CR, DDisc_AUC_200, SSAGA_Educ, DDisc_SV_1yr_200, DDisc_SV_6mo_200, DDisc_SV_3yr_200, LifeSatisf_Unadj, DDisc_SV_5yr_200, ListSort_AgeAdj, ReadEng_Unadj, SCPT_TN, SCPT_SPEC, ReadEng_AgeAdj, ListSort_Unadj, DDisc_AUC_40K, DDisc_SV_10yr_200, DDisc_SV_5yr_40K, PicSeq_AgeAdj, SSAGA_TB_Yrs_Since_Quit, PicSeq_Unadj, DDisc_SV_3yr_40K, DDisc_SV_1yr_40K, SSAGA_Income, Dexterity_AgeAdj, DDisc_SV_10yr_40K, Dexterity_Unadj, DDisc_SV_6mo_40K, DDisc_SV_1mo_200, FamHist_Fath_None, ProcSpeed_AgeAdj, Endurance_AgeAdj, Endurance_Unadj, DDisc_SV_1mo_40K, SSAGA_TB_Age_1st_Cig, ASR_Rule_Pct, ASR_Thot_Raw, EVA_Denom, SSAGA_TB_Still_Smoking, ASR_Thot_Pct, PercStress_Unadj, Taste_AgeAdj, ASR_Rule_Raw, Taste_Unadj, AngAggr_Unadj, Times_Used_Any_Tobacco_Today, PSQI_Score, Avg_Weekend_Cigarettes_7days, Avg_Weekend_Any_Tobacco_7days, Total_Cigarettes_7days, Avg_Weekday_Cigarettes_7days, FamHist_Fath_DrgAlc, Num_Days_Used_Any_Tobacco_7days, Total_Any_Tobacco_7days, Avg_Weekday_Any_Tobacco_7days, SCPT_FP, THC, PMAT24_A_S

List of the 38 subject measures (SMs) investigated by Granziol and Cona (2023), with their unique identifiers as provided by the HCP consortium:

AngHostil_Unadj, LifeSatisf_Unadj, PercReject_Unadj, PercStress_Unadj, SelfEff_Unadj, PSQI_Score, ASR_Witd_T, ASR_Thot_T, DSM_Anxi_T, DSM_Depr_T, NEOFAC_C, NEOFAC_N, NEOFAC_E, PainIntens_RawScore, PainInterf_Tscore, DDisc_AUC_200, DDisc_AUC_40K, PicSeq_AgeAdj, PMAT24_A_CR, VSPLOT_TC, SCPT_SPEC, ListSort_AgeAdj, ER40_CR, ReadEng_AgeAdj, Total_Drinks_7days, Total_Any_Tobacco_7days, SSAGA_Times_Used_Illicits, SSAGA_Mj_Times_Used, AngAggr_Unadj, ASR_Rule_T, ASR_Extn_T, DSM_Antis_T, DSM_Hype_Raw, NEOFAC_A, Flanker_AgeAdj, CardSort_AgeAdj, ProcSpeed_AgeAdj, NEOFAC_O

## Supplementary Figures

**Figure S1.**
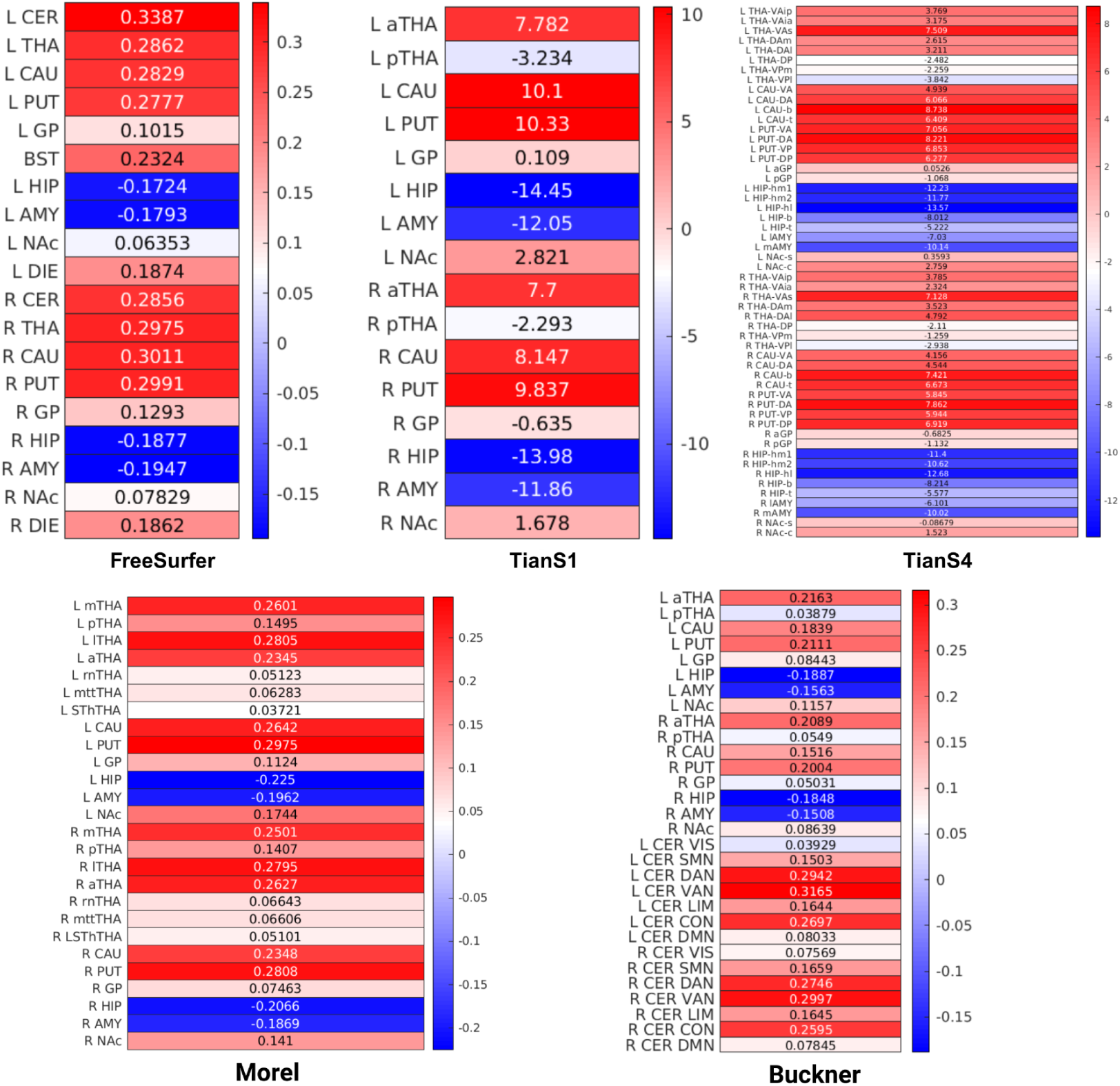
Subcortical clusters for different parcellations. For each subcortical atlas, we plotted the first principal component of the time evolution of the principal eigenvector associated with the sliding-windows temporal correlation, concatenated across subjects. Each of these vectors shows the competitive relationship between two different subcortical clusters for alternative choices of subcortical parcellation **(a)** FreeSurfer subcortical atlas. Labels: CER (cerebellum), THA (thalamus), CAU (caudate nucleus), PUT (putamen), GP (globus pallidus), BST (brainstem), HIP (hippocampus), AMY (amygdala), NAc (nucleus accumbens), DIE (ventral diencephalon). **(b)** Tian S1 parcellation. Labels: same as for FreeSurfer, except aTHA (anterior thalamus), pTHA (posterior thalamus). **(c)** Tian S4 parcellation. Labels: THA-VAip (inferior ventroanterior thalamus, posterior division), THA-VAia (inferior ventroanterior thalamus, anterior division), THA-VAs (superior ventroanterior thalamus), THA-DAm (medial dorsal anterior thalamus), THA-DAl (lateral dorsal anterior thalamus), THA-DP (dorsoposterior thalamus), THA-VPm (medial ventroposterior thalamus), THA-VPl (lateral ventroposterior thalamus), CAU-VA (ventral anterior caudate), CAU-DA (dostal anterior caudate), CAU-b (caudate body), CAU-t (caudate tail), PUT-VA (ventral anterior putamen), PUT-DA (dorsal anterior putamen), PUT-VP (ventral posterior putamen), PUT-DP (dorsal posterior putamen), aGP (anterior globus pallidus), pGP (posterior globus pallidus), HIP-hm1 (hippocampus head medial subdivision 1), HIP-hm2 (hippocampus head medial subdivision 2), HIP-hl (hippocampus head lateral subdivision), HIP-b (hippocampus body), HIP-t (hippocampus tail), lAMY (lateral amygdala), mAMY (medial amygdala), NAc-s (nucleus accumbens shell), NAc-c (nucleus accumbens core) **(d)** Tian S1 + thalamic Morel parcellation. aTHA (anterior thalamus), lTHA (lateral thalamus), mTHA (medial thalamus), pTHA (posterior thalamus), rnTHA (red nucleus), mttTHA (mammillothalamic tract), SThTHA (subthalamic nucleus) **(e)** Tian S1 + cerebellar Bucker parcellation.

**Fig. S2.**
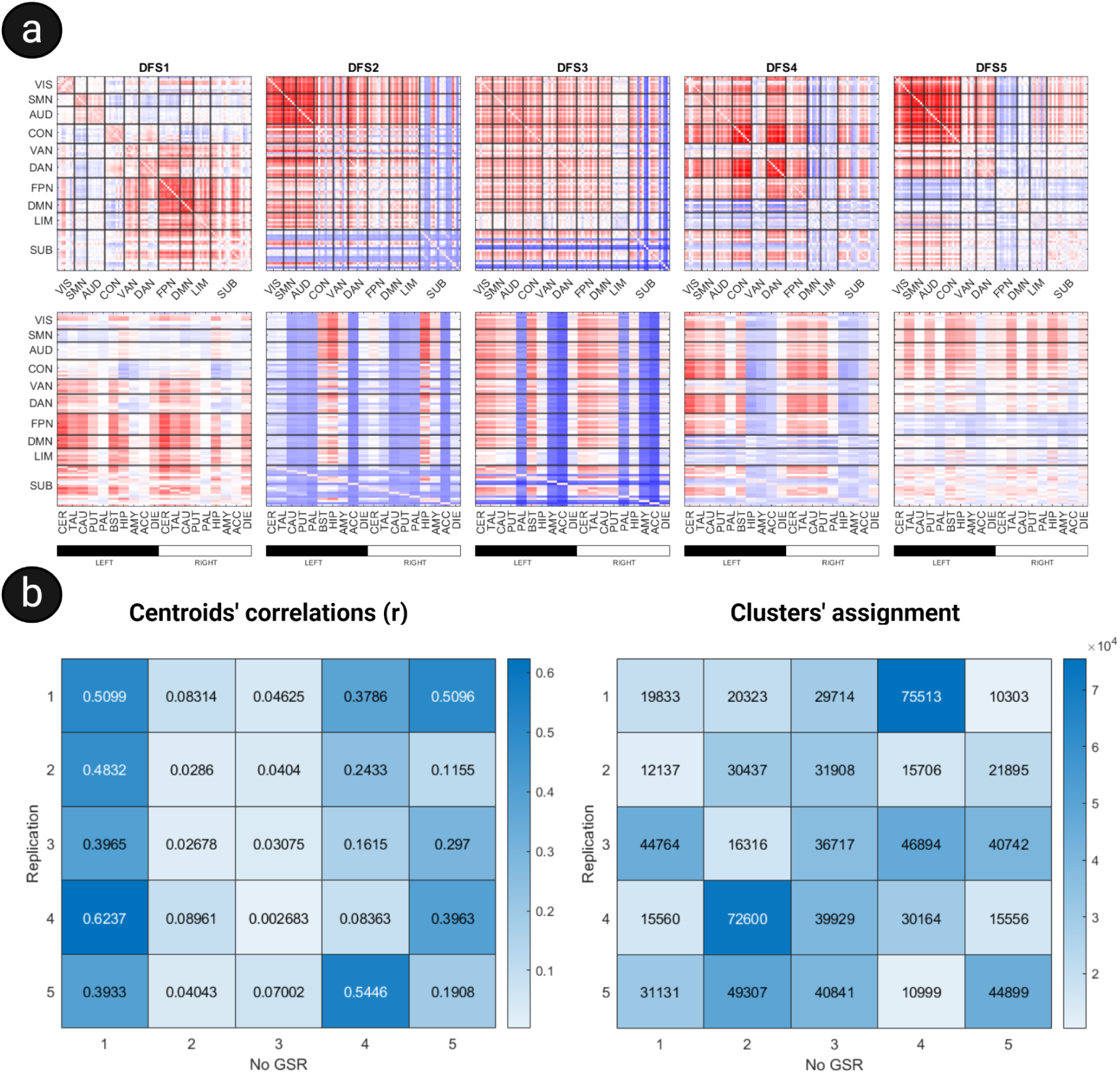
Preprocessing effects: Global Signal Regression. **(a)** Dynamic Functional States obtained without regressing the global signal. Importantly, the aspect of the DFSs qualitatively affected by this methodological choice, which is consistent with the ongoing debate in the literature about GSR. **(b)** This is also quantitatively assessed through these confusion matrices that display correlation values between the centroids (left) and clusters’ assignment with and without GSR (right).

**Fig. S3.**
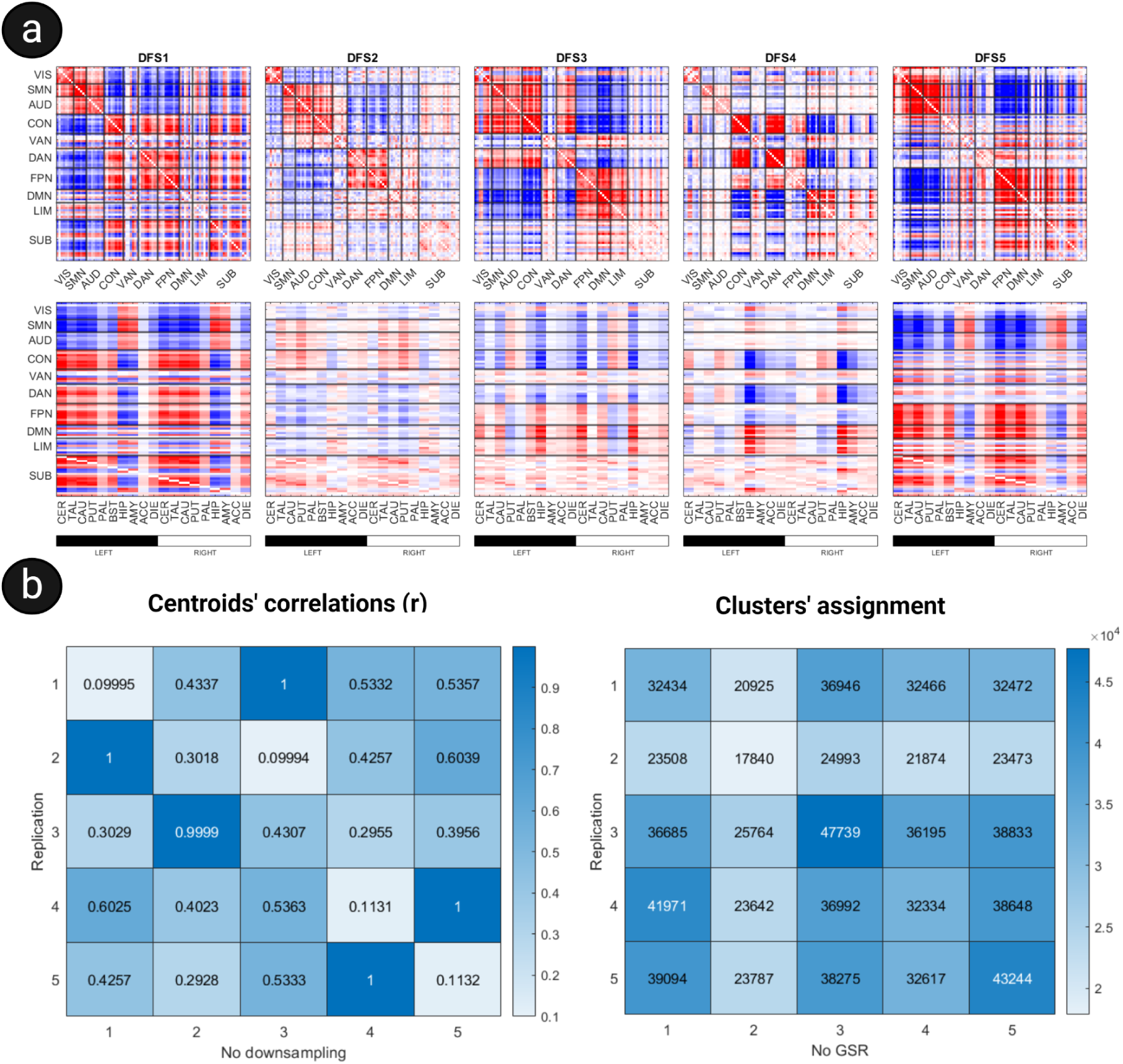
Preprocessing effects: downsampling. **(a)** Dynamic Functional States obtained without downsampling the timeseries. Predictably, the qualitative aspect of the DFSs is not evidently affected by this methodological choice. **(b)** This is also quantitatively assessed through these confusion matrices that display correlation values between the centroids (left) and clusters’ assignment with and without downsampling of the timeseries (right).

**Fig. S4.**
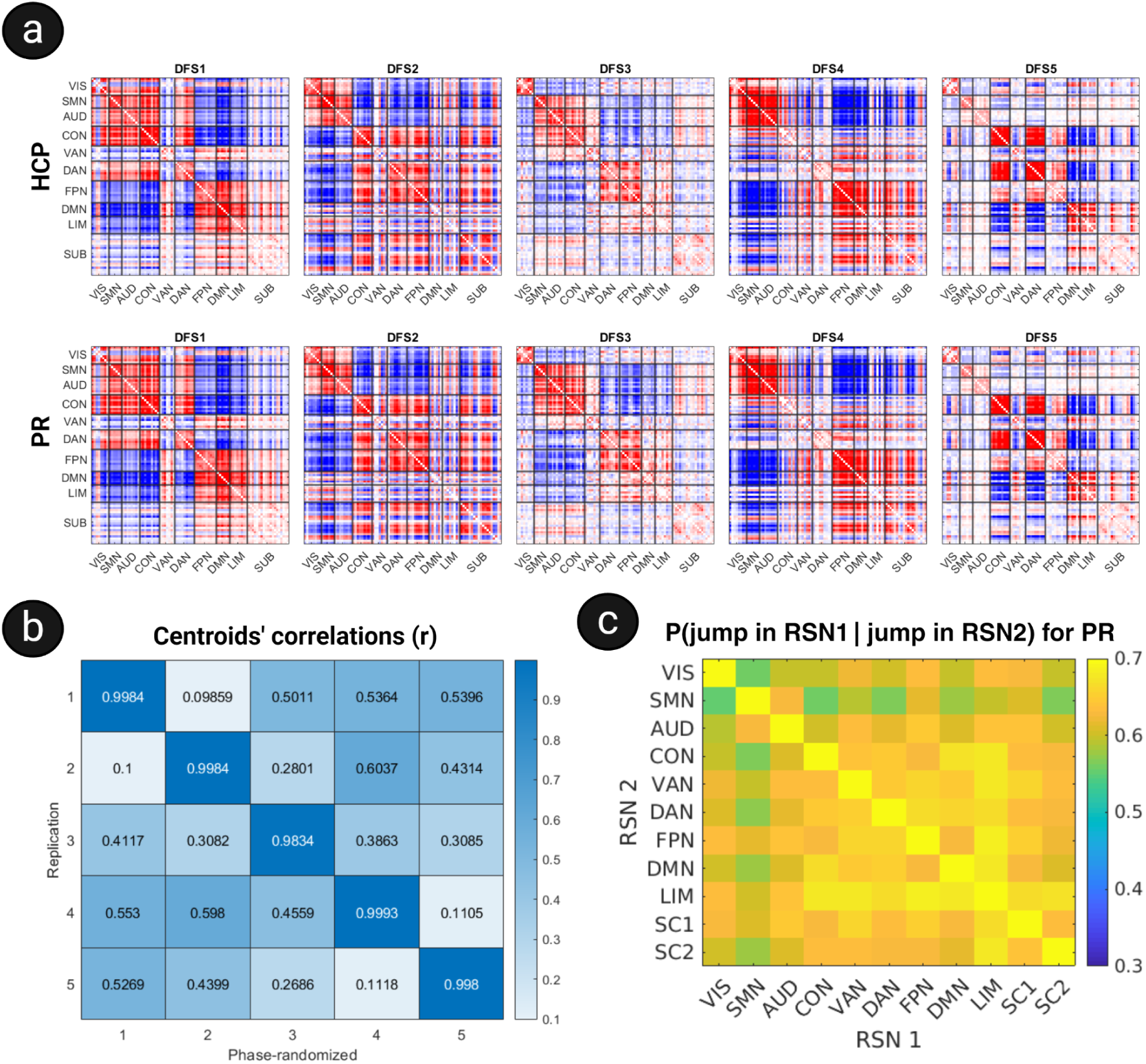
Phase randomization’s results. **(a)** Matrix representation of the cluster centroids of the K=5 dynamic functional states (DFS) found for the original HCP data (top) and for phase-randomized data (bottom). The original and PR results are barely distinguishable . **(b)** A confusion matrix with Pearson correlation values for each couple of DFS centroids between the replication study (y axis) and PR-generated data (x axis). Original and PR centroids match almost perfectly **(c)** Conditional probability matrices where we plot conditional probabilities P(p_i_|p_j_) for each couple of networks, where p_i_ and p_j_ are the probabilities of a connectivity jump in the network i and j respectively. PR data qualitatively maintains the structure of conditional jump probabilities, but conditional probabilities are generally larger (KS test, P<0.05).

**Fig. S5.**
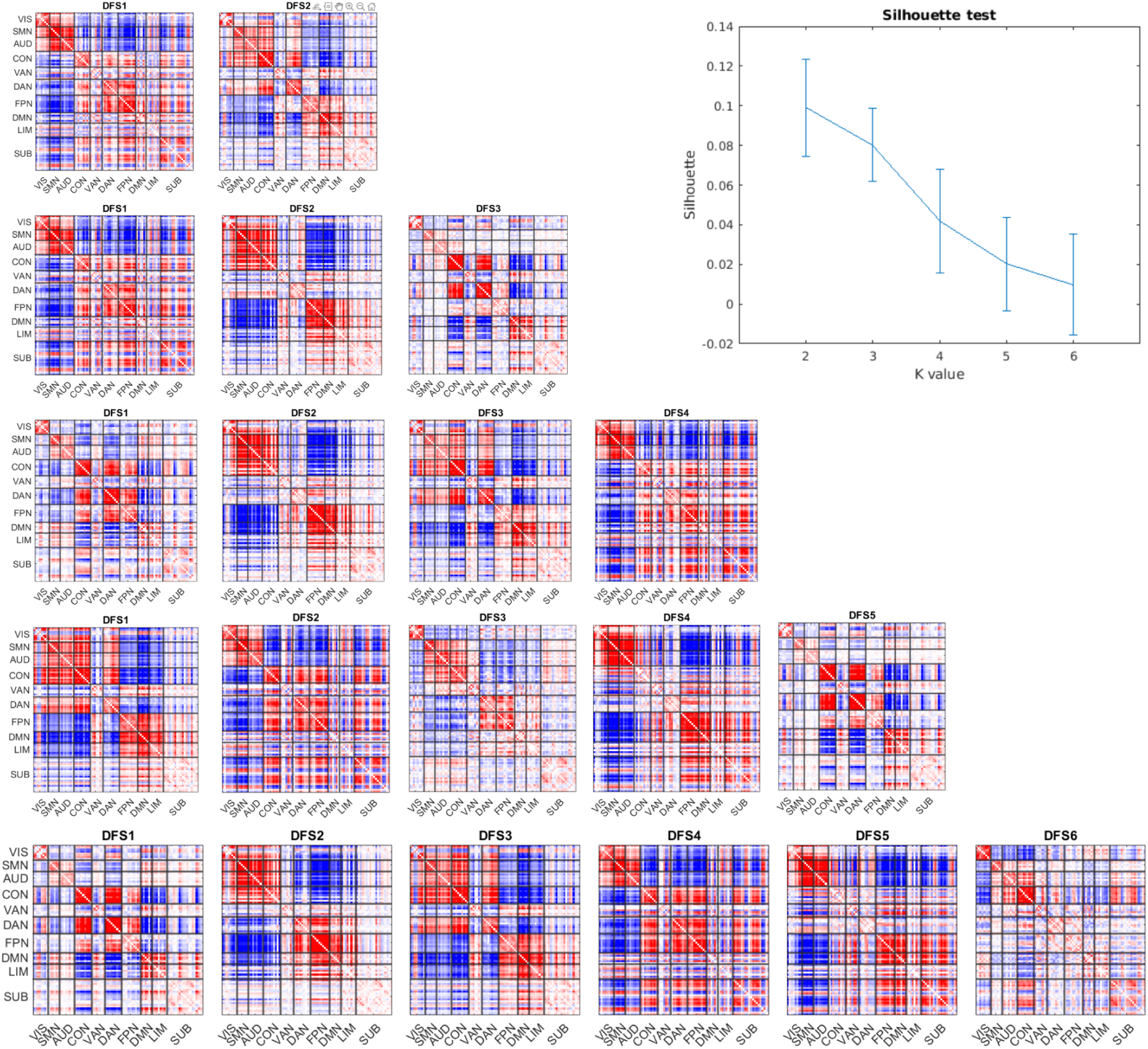
Dynamical Functional States for K = 2, . . ., 6. DFSs obtained for different values of K, from 2 to 6, with the GordonLaumann cortical atlas (and corresponding Silhouette values). As appreciable from this figure, increasing the number of clusters led to the inclusion of new states without altering the original set present for K=2.

**Fig. S6.**
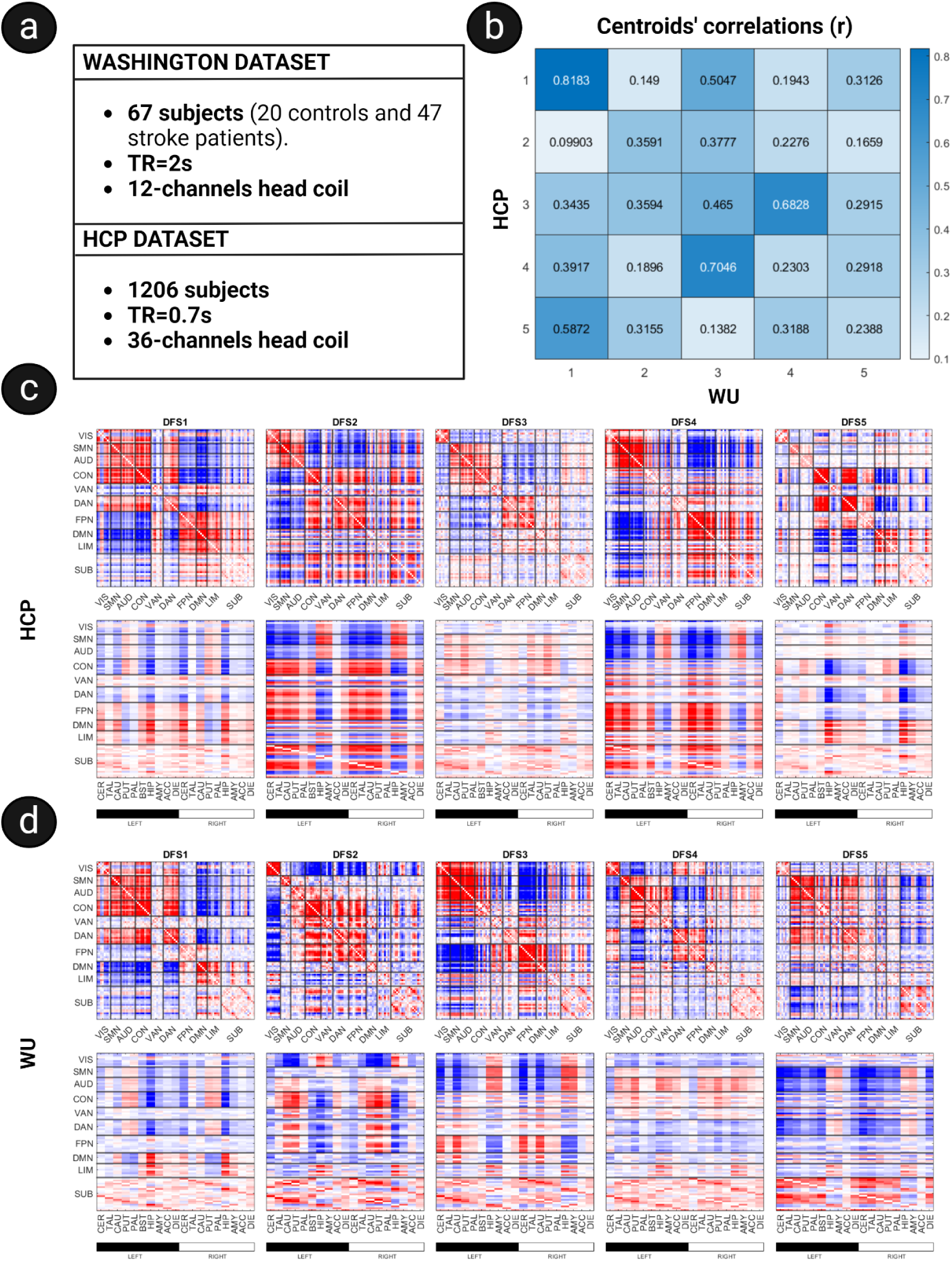
Replication study. **(a)** The main features of the two datasets. **(b)** A confusion matrix with Pearson correlation values for each couple of DFS centroids between the replication study (HCP) and FA22 (WU). **(c)** The cluster centroids of the K=5 dynamic functional states (DFSs) found in the HCP data set. Each centroid *ν* is shown in matrix form, by plotting the matrix *ν* x *ν*^t^. **(d)** As in **(c)**, but for the Washington dataset. The bottom row of **(c)** and **(d)** is a zoom in of the cortico-subcortical interaction.

**Fig. S7.**
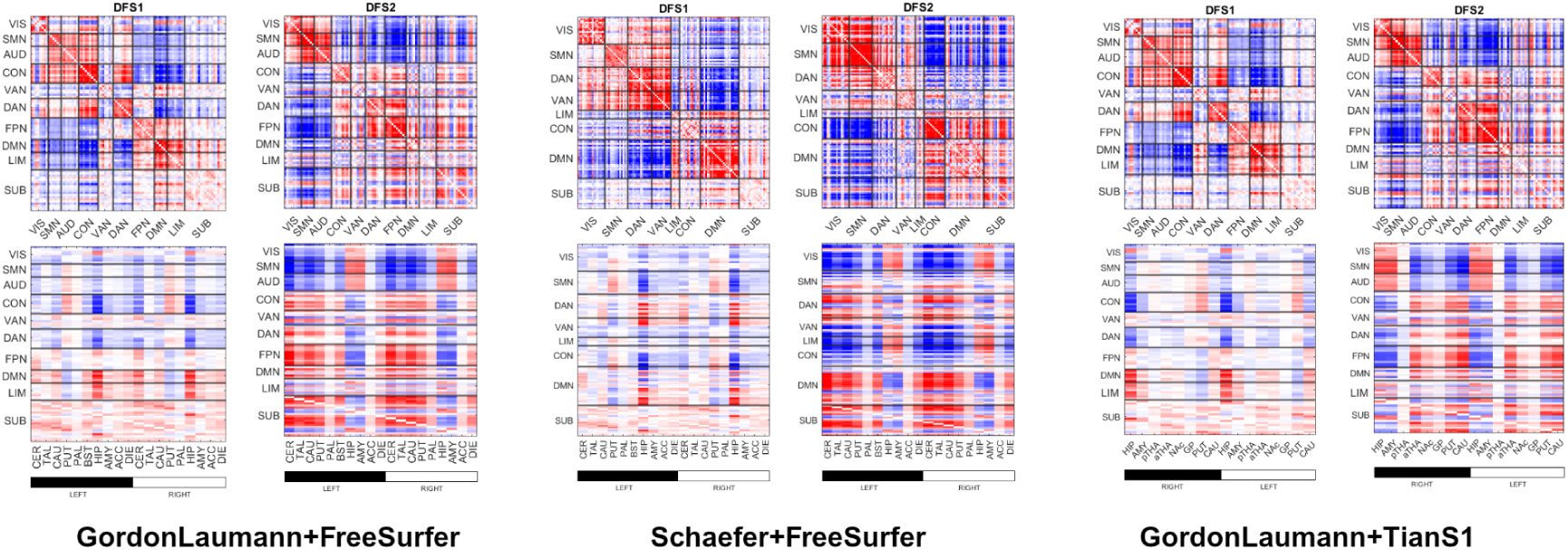
We show the cluster centroids of the K=2 dynamic functional states (DFSs) found in the HCP data set for different choices of cortical and subcortical parcellations. Each centroid *ν* is shown in matrix form, by plotting the matrix *ν* x *ν*^t^.

**Fig. S8.**
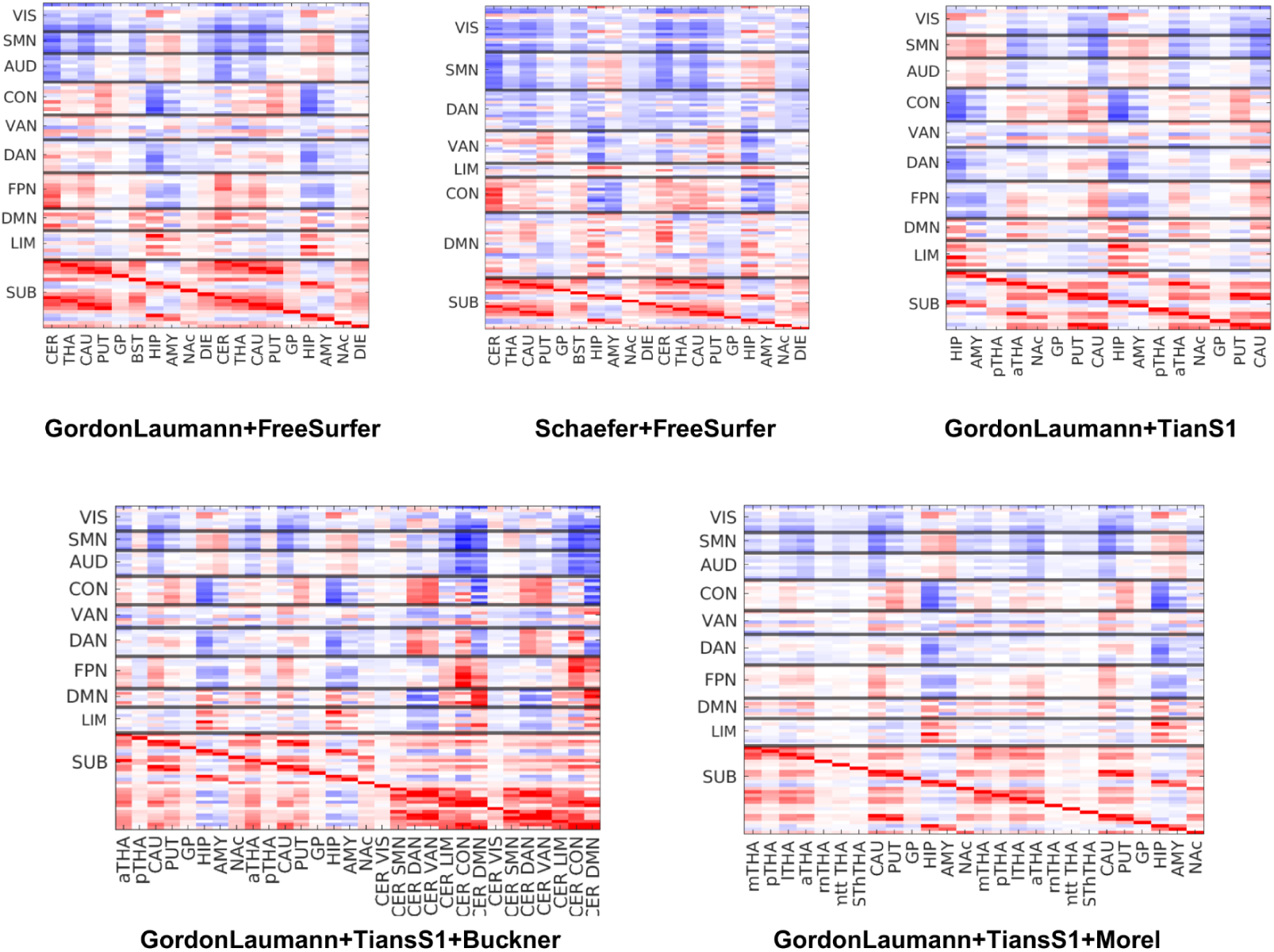
Subcortical static functional connectivity.

